# Evaluation of a *Chlamydia trachomatis*-specific, commercial, real-time PCR for use with ocular swabs

**DOI:** 10.1101/245605

**Authors:** H Pickering, MJ Holland, AR Last, MJ Burton, SE Burr

**Affiliations:** London School of Hygiene and Tropical Medicine, London, UK

**Keywords:** *Chlamydia trachomatis*, diagnostics, real-time PCR, ocular swabs

## Abstract

**Background:** *Chlamydia trachomatis* (*Ct*) is the most common bacterial sexually transmitted infection and the causative organism of trachoma, the leading infectious cause of blindness worldwide. Trachoma is diagnosed clinically by observation of conjunctival inflammation and/or scarring, however, there is evidence that monitoring *Ct* infection may be required for elimination programs. There are many commercial and ‘in-house’ nucleic acid amplification tests for the detection of *Ct* DNA, but the vast majority have not been validated for use with ocular swabs. This study evaluated a commercial assay, the Fast-Track Vaginal swab kit, using conjunctival samples from two trachoma-endemic areas. An objective, biostatistical-based method for binary classification of continuous PCR data was also developed, to limit potential user-bias in diagnostic settings.

**Results:** The Fast-Track Vaginal swab assay was run on 210 ocular swab samples from Guinea-Bissau and Tanzania. Fit of individual amplification curves to exponential or sigmoid models, derivative and second derivative of the curves and final fluorescence value were examined for utility in thresholding for determining positivity. The results from the Fast-Track Vaginal swab assay were evaluated against a commercial test (Amplicor CT/NG) as well as a non-commercial test (in-house ddPCR) both of whose performance has previously been evaluated.

Significant evidence of exponential amplification (R^2^ > 0.99) and final fluorescence > 0.15 were combined for thresholding. This objective approach identified a population of positive samples, however there were a subset of samples that amplified towards the end of the cycling protocol (at or later than 35 cycles), which were less clearly defined. The Fast-Track Vaginal swab assay showed good sensitivity against the commercial (95.71) and non-commercial (97.18) tests. Specificity was lower against both tests (90.00 and 96.55 respectively).

**Conclusions:** This study defined a simple, automated protocol for binary classification of continuous, real time qPCR data, for use in an end-point diagnostic test. This method identified a population of positive samples, however, as with manual thresholding, a subset of samples that amplified towards the end of the thermal cycling program were less easily classified. When used with ocular swabs, the Fast-Track Vaginal swab assay had good sensitivity but for *Ct* detection lower specificity than the commercial and non-commercial assays it was evaluated against, possibly leading to false positives.

## Background

*Chlamydia trachomatis* (*Ct*) is the most common bacterial sexually transmitted infection [1] and the leading infectious cause of blindness worldwide [2]. Trachoma, conjunctival *Ct* infection and subsequent disease, is responsible for visual impairment or blindness in an estimated 2.2 million people [2]. Trachoma is targeted for elimination by 2020 [3], using a series of interventions known as SAFE [4]; Surgery for trichiasis (inturned eyelashes), Antibiotics for infection [community mass drug administration (MDA)], Facial cleanliness and Environmental improvement to reduce transmission. Control programs use clinical diagnosis of trachoma for monitoring prevalence, chiefly signs of trachomatous inflammation-follicular (TF) and trachomatous inflammation-intense (TI). However, the correlation between TF/TI and conjunctival *Ct* infection lessens as prevalence declines [5, [6], suggesting tests for infection may be necessary to accurately monitor trachoma dynamics, particularly in low-endemicity and post-MDA settings [7].

*Ct* infection has historically been diagnosed through culture of the bacterium, antigen detection and direct cytological examination [8]. Currently, nucleic acid amplification tests (NAATs) are the gold standard for *Ct* detection as they are more sensitive and allow increased throughput. Many commercial and non-commercial assays are available, however, the majority have not been validated for use with ocular swabs. Incorporating tests for *Ct* infection into trachoma control programmes, rather than relying on clinical diagnosis alone, has the potential to reduce cost and increase success [9].

Positivity in NAATs based on real time qPCR assays, such as Artus *Ct* Plus RG PCR Kit (Qiagen), is primarily based on samples ‘displaying an exponential trace’ [10] during thermal cycling, and there are a number of methods for binary classification of samples into positives and negatives. It is common for thresholds to be defined based on manual inspection of traces, as well as by comparison to positive and negative controls. Samples that demonstrate exponential amplification before this manually defined threshold are deemed positive. This inherently subjective method introduces user-bias.

This study evaluated a commercial assay, the Fast-Track Vaginal swab kit, for diagnosing *Ct* infection from ocular swabs. Additionally, the study aimed to define an objective method for binary classification of samples using the raw PCR amplification curves, rather than subjective thresholding by individual users.

## Methods

### Sample collection

Samples were collected from the upper tarsal conjunctiva using a Dacron polyester–tipped swab (Hardwood Products Company, Guilford, Maine). The swab was passed firmly four times across the conjunctiva with a quarter turn between each pass. All samples were kept on ice packs in the field until transfer to –80°C the same day for storage until processing. Samples included were collected in Kilimanjaro Region, Northern Tanzania [11] (109) and Bijagos Islands, Guinea-Bissau (101) as part of trachoma surveys.

### DNA extraction and amplification

DNA was extracted from all swabs using the QIAmp DNA mini kit (Qiagen, Crawley, UK). For sample processing using the Fast-track Diagnostic (FTD) Vaginal Swab kit (Fast-Track Diagnostics, Esch-sur-Alzette, Luxembourg), 10ul extracted DNA was amplified in a total reaction volume of 25ul. Positive and negative controls were included in each run. Cycling conditions were as described in the manufacturer’s instructions. Raw fluorescence data was modeled to determine positivity as described below. *C. trachomatis* was detected using the Amplicor CT/NG kit (Roche Molecular Systems, Branchburg, NJ) with previously described modifications [12]. Samples whose absorbance values at 450 nm (A_450_) were ≥0.8 were considered positive while samples less than 0.2 A_450_ were negative. Samples for which the result was equivocal (≥0.2, <0.8) were tested again in duplicate. The sample was only considered positive if the A_450_ of one of the retests was ≥0.8. DNA amplification by droplet digital PCR (ddPCR) was conducted as previously described [13]. A sample was considered positive if the 95% confidence interval of *Ct* plasmid copies per µl did not intersect zero.

### Data analysis

The raw fluorescence data for the FTD Vaginal swab PCR were used for all analyses. Data for Amplicor and ddPCR were analysed as previously described [13, [14]. All analyses were performed using R version 3.3.2. Data were fitted to exponential and sigmoid models using the *qpcR* package. The *mixtools* package was used for mixtures modelling. FTD Vaginal swab PCR was evaluated against the Amplicor CT/NG and ddPCR assays, using the *caret* and *psych* packages to calculate sensitivity, specificity, positive predictive value, negative predictive value and Cohens kappa.

## Results

### Defining threshold for positivity

Two-hundred and ten upper tarsal conjunctival swabs were tested for the presence of *Ct* infection using the FTD Vaginal swab kit. Data from all 210 samples was used to determine an automated threshold by which samples could be considered as either positive or negative.

Raw amplification curves were fitted to exponential and sigmoid models (four-parameter logistic regression) and evaluated using their respective R^2^ values, to test for evidence of exponential amplification. Samples evidenced by a clear exponential trace and final fluorescence intensity, including all positive controls, fitted both models well. However, samples that had a final fluorescence less than or equivalent to the negative controls had variable R^2^ values, with many showing strong fits for one or both models. This discrepancy was demonstrated by poor correlation of fluorescence intensity at cycle 40 and the model R^2^values (exponential model; correlation = 0.31, p-value = 0.0001 and sigmoid model; correlation = 0.35, p-value < 0.0001). Good model fit in samples that did not amplify was likely caused by instability in low-level/background fluorescence, leading to fluctuations that can be mistaken for amplification (Figure 1). Using model fit alone would classify these non-amplifying samples as positive, dramatically reducing specificity. Similar results were found with the derivative and second derivative of the sigmoid models.

**Figure 1.**
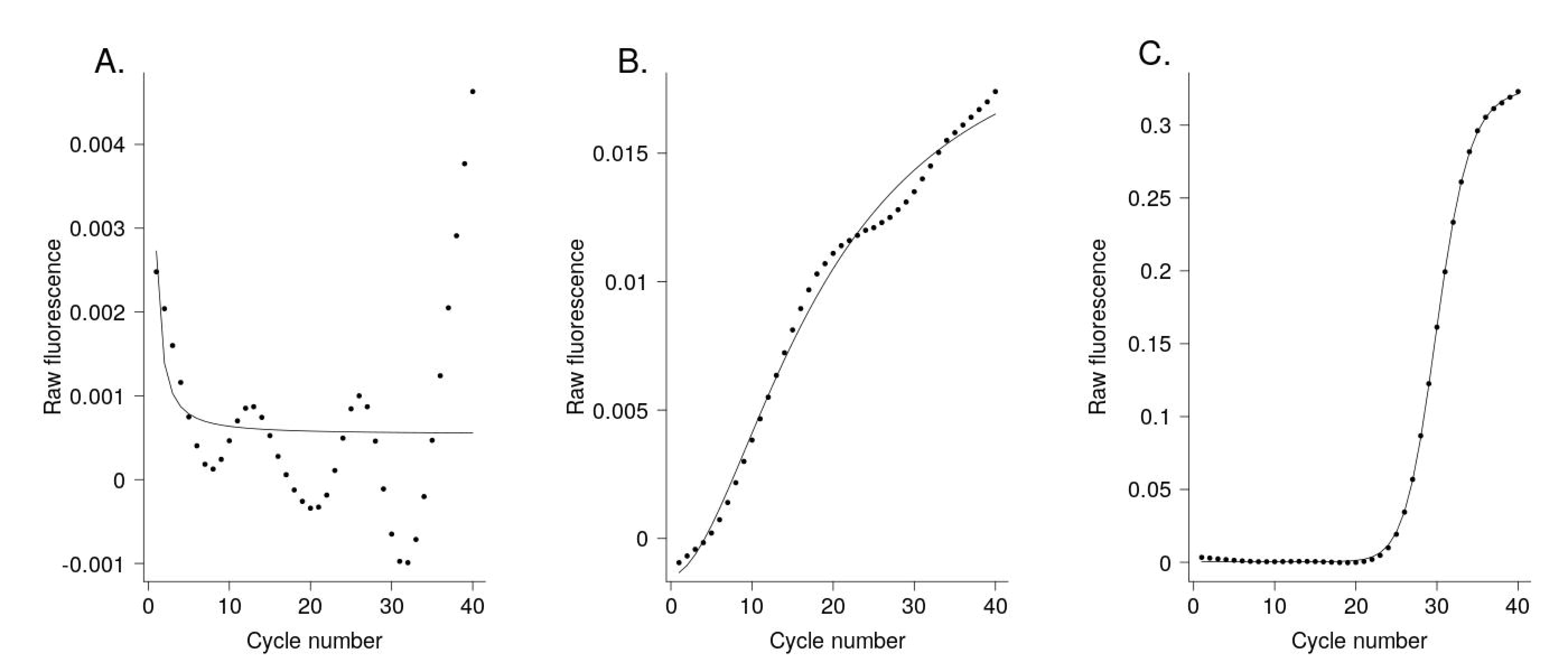
Sigmoid models representative of variation in amplification. Raw fluorescence results from samples processed using the FTD Vaginal swab kit. Sigmoid models representing; (A) samples that did not amplify and do not fit a sigmoid model, (B) samples that did not amplify but do fit a sigmoid model (C) and samples that did amplify and do fit a sigmoid model.

To overcome the limitations of only considering ‘clear’ exponential amplification, a minimum final fluorescence component was incorporated into the method for defining positive samples. The final fluorescence values for each sample were modelled onto mixtures of two or three normal distributions. Allowing three distributions provided the optimum fit (log-likelihood = 474) compared with two distributions (log-likelihood = 431). Both models identified samples with a final fluorescence above 0.15 units as a separate population. A mixture of three distributions also highlighted a population of samples with a final fluorescence between 0.05 and 0.15 units (Figure 2).

**Figure 2.**
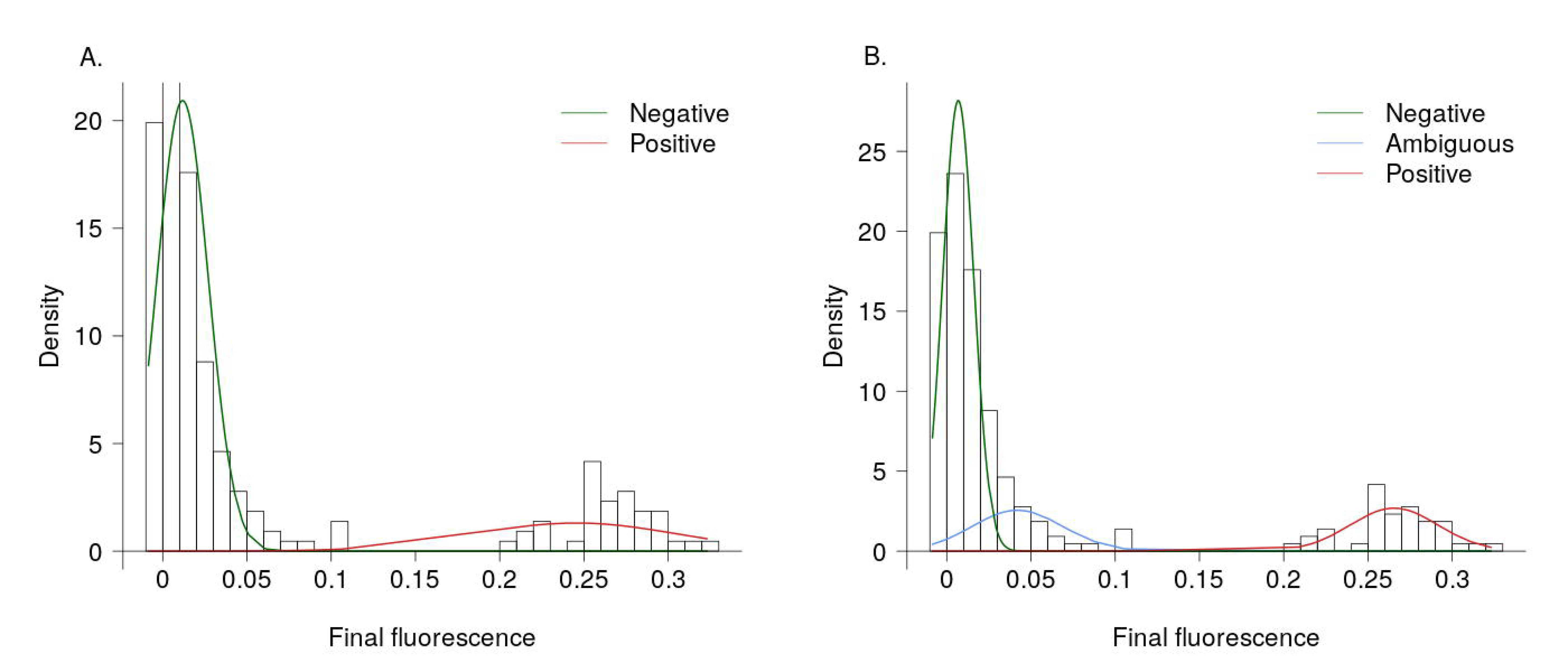
Mixtures models of final fluorescence values. Final fluorescence values, from samples processed using the FTD Vaginal swab kit, were modelled onto mixtures of two (A) and three (B) normal distributions.

### Diagnostic evaluation of FTD Vaginal swab PCR

Finally, we conducted a diagnostic evaluation of the FTD Vaginal swab kit against previously validated commercial (Amplicor CT/NG [14]) and in-house (ddPCR [13]) assays (Table 1), using a subset of 100 samples. For diagnostic evaluation, two methods for identifying exponential amplification were combined, sigmoid model R^2^ value > 0.99 and final fluorescence > 0.15. Thirty of 100 samples were positive by Amplicor CT/NG; the FTD Vaginal swab kit and ddPCR correctly identified 27 of these, respectively calling 3 and 2 additional positive results. Twenty-nine of 100 samples were positive by ddPCR, the FTD Vaginal swab kit correctly identified 28 of these, additionally calling 2 further positive results. In-house ddPCR had greater sensitivity (97.14) compared with Amplicor CT/NG than previously reported [13].

**Table 1.** Diagnostic comparison of FTD-Vaginal swab PCR to Amplicor CT/NG and in-house ddPCR.

|  | **FTD Vaginal swab PCR versus** |  |
| --- | --- | --- |
|  | **Amplicor CT/NG**<br>**(Roche)** | **In-house ddPCR**<br>**(Bio-Rad QX100)** |
| Sensitivity | 95.71 (91.74-99.68) | 97.18 (93.94-100.0) |
| Specificity | 90.00 (84.12-95.88) | 96.55 (92.97-100.0) |
| PPV | 95.71 (91.74-99.68) | 98.57 (96.24-100.0) |
| NPV | 90.00 (84.12-95.88) | 93.33 (88.44-98.22) |
| Cohens Kappa | 0.86 | 0.93 |

*The 95% confidence intervals are shown in brackets. ddPCR, droplet digital PCR. NPV, negative predictive value. PPV, positive predictive value.*

The 3 Amplicor CT/NG-positive samples called as negative by the FTD Vaginal swab kit and by ddPCR were originally identified as equivocal samples before retesting, per the Amplicor protocol. Two of three had no positive droplets by ddPCR and a final fluorescence by the FTD Vaginal swab kit of less than 0.01. The remaining sample did not have enough positive droplets to be reliably called as positive by ddPCR, but had a final fluorescence above 0.1 by the FTD Vaginal swab kit.

## Discussion

This study evaluated results for detection of *Ct* DNA from ocular swabs using the FTD Vaginal swab kit, a commercial PCR assay validated for use with urogenital samples, against previously validated commercial and in-house assays. Automated methods for determining thresholds for positivity from raw amplification curves were also explored. A composite of amplification curve fit and absolute level of amplification was determined to be the best method for the identification of positive results, although, as with setting a manual threshold, there was some ambiguity assigning samples that amplified at or later than 35 cycles. The FTD Vaginal swab kit performed well against both comparator assays, however specificity was notably lower versus Amplicor CT/NG.

Manual and inherently subjective binary classification of continuous real time qPCR data is a significant problem, as it creates unnecessary variability within and across assays. Visual inspection of curves focuses on identifying clear evidence of exponential/sigmoid amplification. This study found that observation of exponential amplification could not singularly define positivity, due to instability in low-level/background fluorescence being mistaken for true amplification. Including mixtures-model clusters defined using the final fluorescence value greatly improved identification of positive samples. For diagnostic evaluation, a sample was deemed positive if; 1) the amplification curve was fit strongly to a sigmoid model (R^2^value > 0.99) and 2) the sample clustered within the right-most population of a mixtures-model of three normal distributions (final fluorescence > 0.15). This objective method identified a population of positive samples, however there was some ambiguity in samples that appear to amplify late in the reaction (at or later than 35 cycles), a problem common to manual threshold setting and inspection. A subset of samples, which formed the middle population of the three distribution mixtures-model, described above, showed varying levels of amplification using the FTD Vaginal swab assay, with final fluorescence values between 0.05 and 0.15. Of interest, the sample in this subset with the highest final fluorescence was positive by Amplicor CT/NG, after an initial equivocal result. As with Amplicor CT/NG, samples that fit in this middle, less clear population should, ideally, be retested.

The FTD Vaginal swab assay performed well against both Amplicor CT/NG and ddPCR. Sensitivity was above 95% for both, with 3 and 1 false negatives respectively. One of the false negatives, was a sample with a final fluorescence over 0.1, which was initially equivocal by Amplicor CT/NG. It is possible with retesting, as suggested above, this sample might have been positive. Specificity was slightly lower against both tests, notably down to 90% for Amplicor CT/NG, with 3 and 2 false positives respectively. Compared with either assay, the specificity of FTD-Vaginal swab assay is below that of various alternative NAATs for *Ct* [15].

## Conclusion

Automated, unbiased classification of continuous real time qPCR data into binary results, for diagnostic purposes, is achievable using a simple set of biostatistical rules. The method described allows objective classification of results from qPCR, using the raw output from thermal cycling programs. FTD Vaginal swab PCR for use with ocular swabs has lower specificity than both Amplicor CT/NG and ddPCR for *Ct* detection, challenging its diagnostic utility with this sample type.

## Declarations

### Ethics approval and consent to participate

Samples were collected from trachoma-endemic communities in Tanzania and Guinea-Bissau following ethical approval from the Kilimanjaro Christian Medical Centre (Tanzania), the National Institute for Medical Research in Tanzania, Comitê Nacional de Ética e Saúde in Guinea Bissau and London School of Hygiene and Tropical Medicine (UK). Written, informed consent was obtained from all participants prior to sample collection. In the case of children, consent was obtained from the parent or guardian.

### Consent for publication

Not applicable.

### Availability of data and materials

The datasets generated and analysed during the current study are available from the corresponding author on reasonable request.

## Author contributions

Study design: MJH, SEB

Sample collection: ARL, MJB

Sample processing: SEB

Data analysis: HP, MJH, SEB

Manuscript preparation: HP, MJH, SEB

Manuscript approval: HP, ARL, MJB, MJH, SEB

All authors read and approved the final version of the manuscript.

## Acknowledgements

The authors would like to acknowledge individuals in Guinea-Bissau and Tanzania who provided samples for this study and the local field teams involved in sample collection.

